# Vaccination generates functional progenitor tumor-specific CD8 T cells and long-term tumor control

**DOI:** 10.1101/2024.02.26.582064

**Authors:** Carlos R. Detrés Román, Michael W. Rudloff, Frank Revetta, Natalie R. Favret, Kristen A. Murray, Jessica J. Roetman, Megan M. Erwin, Mary K. Washington, Mary Philip

## Abstract

**Background:** Immune checkpoint blockade (ICB) therapies are an important treatment for patients with advanced cancers; however only a subset of patients with certain types of cancer achieves durable remissions. Cancer vaccines are an attractive strategy to boost patient immune responses, but less is known about whether and how immunization can induce long-term tumor immune reprogramming and arrest cancer progression. We developed a clinically-relevant genetic cancer mouse model in which hepatocytes sporadically undergo oncogenic transformation. We compared how tumor-specific CD8 T cells (TST) differentiate in mice with early sporadic lesions as compared to late lesions and tested how immunotherapeutic strategies, including vaccination and ICB, reprogram TST and impact liver cancer progression.

**Methods:** Mice with a germline floxed allele of the SV40 large T antigen (TAG) undergo spontaneous recombination and activation of the TAG oncogene, leading to rare early pre-cancerous lesions that inevitably progress to established liver cancer. We assessed the immunophenotype and function of TAG-specific CD8 T cells in mice with early and late liver lesions. We vaccinated mice, either alone or in combination with ICB, to test whether these immunotherapeutic interventions could stop liver cancer progression.

**Results:** In mice with early lesions, a subset of TST were PD1^+^ TCF1^+^ TOX^-^ and could produce IFNγ, while TST present in mice with late liver cancers were PD1^+^ TCF1^lo/-^ TOX^+^ and unable to make effector cytokines. Strikingly, vaccination with attenuated TAG epitope-expressing *Listeria monocytogenes* (LM_TAG_) blocked liver cancer development and led to a population of TST that were TCF1^+^ TOX^-^ TST and polyfunctional cytokine producers. In contrast, ICB administration did not slow cancer progression or improve LM_TAG_ vaccine efficacy.

**Conclusion:** Vaccination, but not ICB, generated a population of progenitor TST and halted cancer progression in a clinically relevant model of sporadic liver cancer. In patients with early cancers or at high-risk of cancer recurrence, immunization may be the most effective strategy.

**What is already known on this topic:** Immunotherapy, including immune checkpoint blockade and cancer vaccines, fails to induce long-term remissions in most patients with cancer.

**What this study adds:** Hosts with early lesions but not hosts with advanced cancer retain a progenitor TCF1+ TST population. This population can be reprogrammed and therapeutically exploited by vaccination, but not ICB, to block tumor progression.

**How this study might affect research, practice, or policy:** For people at high-risk of cancer progression, vaccination administered when a responsive progenitor TST population is present may be the optimal immunotherapy to induce long-lasting progression-free survival.

## INTRODUCTION

Immunotherapies such as immune checkpoint blockade (ICB) have reshaped the cancer treatment landscape, inducing long-term remissions in a subset of patients^1^. In contrast, while vaccines have been enormously successful in combating infectious disease, vaccines for non-viral cancers have had more limited success^2^. Most preclinical and clinical studies on tumor-specific CD8 T cell (TST) vaccine responses have been done in the context of established/late tumors^3^. Less is known about how TST respond and differentiate in response to immunotherapy during early stages of tumorigenesis, when few cells harboring oncogenic mutations and potential neoantigens are present^4^.

We previously developed an autochthonous mouse model of liver cancer (AST;Cre-ER^T2^) in which we could initiate liver carcinogenesis with tamoxifen (TAM)-induced Cre-mediated SV40 large T antigen (TAG) expression in hepatocytes^5^. TAG acts both as an oncogene, driving liver carcinogenesis, and as a tumor-specific neoantigen recognized by CD8 T cells; this model allows precise temporal control of the duration of TST interactions with transformed hepatocytes and tumors. However, in contrast to human tumors, which arise sporadically and progress clonally^4^, TAM-induced oncogene induction is highly efficient, resulting in high antigen burden even in early-stage lesions. To better model sporadic cancer formation and subsequent progression, we allowed AST;Cre-ER^T2^ mice to undergo stochastic TAG oncogene activation through sporadic, TAM-independent Cre-mediated activity.

We found that TST in AST;Cre-ER^T2^ mice with early lesions upregulated PD1 and proliferated. Progenitor PD1+ TST which express the transcription factor (TF) TCF1 and are localized to secondary lymphoid organs or tertiary lymphoid structures have been shown to maintain the TST population and mediate immunotherapy responses^6–9^ and reviewed in^10^ ^11^. In mice with early lesions, we found that a subset of PD1^+^ TST expressed the transcription factor TCF1 and retained the ability to produce IFNγ. This subset was not sustained in mice with late lesions. Here, we tested whether immunization and/or ICB could improve TST function and found that immunization but not ICB generated a robust population of functional TST, which was able to halt prevent cancer progression long-term in liver cancer-prone mice.

## MATERIALS AND METHODS

### Mice

All mice were bred and maintained in a specific pathogen free barrier facility at Vanderbilt University Medical Center. TCR_TAG_ transgenic mice (Strain 005236), Cre-ER^T^^2^ (Strain 008463), C57BL/6 (Strain 000664), and C57BL/6J Thy1.1 mice (Strain 000406) were purchased from The Jackson Laboratory. TCR_TAG_ mice were crossed to Thy1.1 mice to generate TCR_TAG_;Thy1.1 and TCR_TAG_;Thy1.1/Thy1.2 (Thy1.12) mice. AST [(<u>A</u>lbumin-flox<u>S</u>top-SV40 large <u>T</u> antigen] mice, obtained from Drs. Natalio Garbi and Günter Hämmerling, German Cancer Research Center and previously described^12^, were crossed to Cre-ER^T2^ mice to generate AST;Cre-ER^T2^.

### Tumor cohort setup

Age- and sex-matched AST;Cre-ER^T2^ were assigned to experimental tumor cohorts at 6 weeks age. Both male and female mice were used for experimental cohorts. All tumor cohort mice were monitored twice weekly for body condition score (BCS) and abdominal distension (scored from 1-4: 1 no swelling, 2 mild abdominal enlargement, 3 moderate abdominal enlargement, 4 marked abdominal enlargement). Mice were analyzed at endpoint or specified time points as specified in figure legends and text. For survival studies, mice were euthanized at pre-specified endpoints: (BCS 2), abdominal girth score 4, or difficulty with movement or eating/drinking. For intervention studies (LM vaccination and ICB), mice in AST;Cre-ER^T2^ tumor cohorts were randomly assigned to treatment arms. Sample size was determined to obtain 80% power to detect a 35% difference between groups, based on our previously observed/published differences and reproducibility. Potential confounders such as animal/cage location, treatment order, and measurement order were minimized by co-housing mice receiving different treatments. CRDR and MP were aware of the group allocation throughout experiments.

### Liver tissue preparation, fixing and paraffin block

Liver tissue was removed from AST;Cre-ER^T2^ mice and immediately transferred into 10% buffered formalin for a minimum of 48 hours before transfer and storage in 70% ethanol. The Vanderbilt Translational Pathology Shared Resource (TPSR) performed paraffin embedding as follows: 70% ethanol for 30 minutes, 80% ethanol for 45 minutes, 95% ethanol for 45 minutes (twice), 100% ethanol for 45 minutes, 100% ethanol for 35 minutes (twice), xylene for 35 minutes (twice), wax (paraffin) for 60 minutes (3 times).

### Immunohistochemistry

SV40 TAG-specific mouse monoclonal antibody (BD Bioscience 554149) primary antibody was used. Antigen Retrieval was performed with pH 6.0 citrate buffer at 105°C pressure cooker for 15 minutes and a 10 minute bench cool down. Mouse on Mouse (MOM) Ig Block was performed with Vector MKB-2213 and incubated for 60 minutes. Peroxidase Block was performed at 0.03% H202 w/sodium azide and incubated for 5 minutes. Primary antibody was diluted at 1:800 and incubated for 60 minutes. Detection was performed with Dako EnVision+ System-HRP Labeled Polymer and incubated for 30 minutes. Chromogen was performed with DAB+ and incubated for 5 minutes.

### Cell isolation

Spleens were mechanically disrupted to a single-cell suspension with the back of a 3 mL syringe plunger, passed through a 70 μm strainer, and lysed with ammonium chloride potassium (ACK) buffer (150 mmol/L NH_4_Cl, 10 mmol/L KHCO_3_, 0.1 mmol/L Na_2_EDTA). Cells were washed once and resuspended with RPMI-10: RPMI 1640 (Corning MT10040CV) supplemented with 10% FBS (Corning 35010CV). Livers were mechanically disrupted to a single-cell suspension using a glass pestle against a 150 μm metal mesh in cold PBS containing 2% FBS (2% FBS) and filtered through a 100 μm strainer. The liver homogenate was spun down at 400g for 5 min at 4°C, and the pellet was resuspended in 15 mL 2% FBS, 500 U heparin (NDC 63323– 540–05), and 10 mL PBS Buffered Percoll (Cytiva 17089102), mixed by inversion, and spun at 500 g for 10 min at 4°C. Pellets, enriched for liver-infiltrating lymphocytes, were lysed with ACK buffer, and cells were resuspended in RPMI-10 for downstream analysis. Periportal and celiac lymph nodes were collected and pooled for liver-draining lymph node (ldLN) analysis. Lymph nodes were mechanically dissociated into single-cell solutions using the textured surface of two frosted microscope slides into cold RPMI-10. Isolated leukocytes from spleen, lymph nodes, and liver were analyzed by flow cytometry as described below.

### Adoptive T cell transfer

To transfer naive TCR_TAG_ T cells into AST;Cre-ER^T2^mice, splenocytes were sterilely-isolated and processed from TCR_TAG_;Thy1.1 or TCR_TAG_;Thy1.12 transgenic mice as described above. Next, an aliquot was taken from the single-cell suspension and counted via hemocytometer to obtain total splenocyte concentration, stained with antibodies for CD8, Thy1.1 and Thy1.12 to determine the percentage of CD8+ Thy1.1+ or Thy1.12+ TCR_TAG_. These cells were resuspended in RPMI and injected i.v. into each recipient mouse. For the experiments in Figures 2-3, 2.5×10^6^ CD8+ TCR_TAG_;Thy1.1 and/or TCR_TAG_;Thy1.12 were injected per mouse. For experiments in Figures 4-5 0.5×10^6^ TCR_TAG_;Thy1.1/mouse were injected.

### CFSE labeling

Splenocytes were isolated and processed as described above and resuspended in 2.5 mL of RPMI. They were then rapidly mixed with equal volume of 2x CFSE [10 μM] solution and incubated for 5 min at 37°C at a final CFSE concentration [5 μM]. Stained cells were quenched by mixing with 5 ml pure FBS, washed twice with RPMI, and resuspended in RPMI for injection.

### *Listeria* infection

*Listeria monocytogenes* (LM) ΔactA ΔinlB expressing Tag-I epitope (SAINNYAQKL, SV40 large T antigen_206–215_) (LM_TAG_)^5^, or without exogenous antigens (LMΦ), were generated by Aduro Biotech and stored at −80°C. Mice were infected with 5×10^6^ c.f.u. of LM_TAG_ or LMΦ via i.v. injection.

### Immune Checkpoint Blockade

Anti-PD1 (clone RMP1-14) and anti-PDL1 (clone 10.F.9G2) antibodies or isotype control (clone LTA-2) were purchased from BioXcell. Antibodies were diluted in 1X sterile PBS and injected i.p. every other day for 5 doses, at 200 µg per antibody per mouse.

### Antibodies and reagents

Fluorochrome-conjugated antibodies and cell dyes were purchased from Miltenyi Biotech, Thermo, BioLegend, Tonbo/Cytek Biosciences, and Cell Signaling Technology. Specific antibodies are listed in Table 1, and cell dyes are listed in Table 2.

### Cell-surface and intracellular cytokine staining

Splenocytes or liver-infiltrating lymphocytes from naive TCR_TAG_ mice or AST;Cre-ER^T^^2^ mice were stained with Ghost Dye Red 780 Viability Dye (1:2,000 dilution) Tonbo/Cytek 13–0865-T500) and antibodies against surface molecules (CD8, Thy1.1, Thy1.12, CD44, PD1) in 2% FBS. For detailed antibody information see Supplementary Table S1 and for cell dye information see Supplementary Table S2. Flow cytometry plots shown in figures are gated on live CD8+ Thy1.1+ or Thy1.12+ transgenic TCR_TAG_ cells. The gating strategy is shown in **Supplementary Fig. S1**. For intracellular staining, splenocytes or liver-infiltrating lymphocytes were surface stained as above and then fixed and permeabilized using the FoxP3 Transcription Factor Fix/Perm (Tonbo/Cytek TNB-0607) per the manufacturer’s instructions before staining for intracellular molecules (TCF1, TOX). The samples were then analyzed by flow cytometry (see Flow cytometric analysis). For analysis of effector cytokine production (IFNγ, TNFα), ex vivo peptide stimulation was performed prior to staining. Splenocytes from naive or tumor-bearing AST;Cre-ER^T2^ mice were mixed with 2×10^6^ C57BL/6 splenocytes and incubated in RPMI-10 for 4 hours at 37°C in the presence of brefeldin A (Biolegend 420601) and TAG peptide (SAINNYAQKL [0.5 μM]; Genscript; custom-synthesized). The cells were then surface-stained (CD8, Thy1.1, Thy1.12), fixed and permeabilized as described above, stained with antibodies against IFNγ and TNFα and analyzed by flow cytometry.

### Flow cytometric analysis

Flow cytometric analysis was performed using an Attune NxT 4 laser Acoustic Focusing Cytometer (ThermoFisher). Flow data were analyzed with FlowJo v10 software (BD Biosciences).

### Statistics

Survival curves were generated using the Kaplan-Meier method and compared using the log-rank (Mantel-Cox) test. For comparison of multiple survival curves, significance threshold was set using Bonferroni correction. For comparisons between two groups, Student t-test was performed. For multiple comparisons, one-way or two-way ANOVA was performed, followed by either post-hoc Tukey or Šidák test. The alpha level was set at 0.05. GraphPad Prism 10 was used for all statistical analyses.

## RESULTS

### AST;Cre-ER^T^^2^ mice sporadically develop liver tumors which progress with age

We previously developed an autochthonous mouse model of liver cancer (AST;Cre-ER^T^^2^) in which we can study TST interactions with cancer cells throughout carcinogenesis^5^. In AST;Cre-ER^T2^ mice, activation of Cre recombinase by a single dose of tamoxifen (TAM) induces expression of the SV40 large T antigen (TAG) under control of the albumin promoter/enhancer in hepatocytes. AST;Cre-ER^T2^ develop large liver tumors within 60-80d post-TAM. To study tumor-specific CD8 T cell (TST) responses against TAG-driven tumors, we used congenic donor lymphocytes from transgenic mice in which CD8 T cells express a single T cell receptor (TCR) specific for TAG epitope-I (TCR_TAG_)^13^. We found that TST became dysfunctional in TAM-treated AST;Cre-ER^T^^2^ mice and were unable to halt tumor progression^5^. However, TAM-treated AST;Cre-ER^T2^ mice had a substantial tumor antigen burden, even at early-stages of tumorigenesis, due to efficient TAM-induced Cre recombination. In most patients, oncogene activation occurs sporadically, thus we sought to determine how TST would differentiate and function in mice with rare sporadic transformed hepatocytes or early liver lesions.

In Cre-ER^T2^ mice, Cre recombinase can undergo stochastic tamoxifen-independent nuclear translocation^14^ ^15^, putting hepatocytes in AST;Cre-ER^T2^ mice at risk of spontaneous TAG oncogene activation. Indeed, we found that as AST;Cre-ER^T2^ mice aged, they reproducibly developed liver tumors (**Fig. 1A**) and progressed to endpoint within ∼225 days (d) (**Fig. 1B**), much later than TAM-induced liver carcinogenesis. Liver tumors become grossly visible only after AST;Cre-ER^T2^ mice were >90d old, and mice >130d old had large tumors (**Fig. 1A**). For subsequent studies, we grouped 40-80d aged mice without macroscopic liver lesions as “early”, 90-120d age mice with small visible tumors as “intermediate”, and >130d aged mice with large tumors as “late” liver lesion time points (**Fig. 1A**). We performed immunohistochemistry staining for TAG on livers from early, intermediate, and late AST;Cre-ER^T2^ mice. While early AST;Cre-ER^T2^ mice had rare small foci of TAG-positive hepatocytes, intermediate and late mice showed progressively larger TAG-positive lesions (**Fig. 1A**). The liver weight of AST;Cre-ER^T2^ mice increased due to tumor burden, with an initial slow growth phase followed by a more rapid growth phase (**Fig. 1C**).

**Figure 1:**
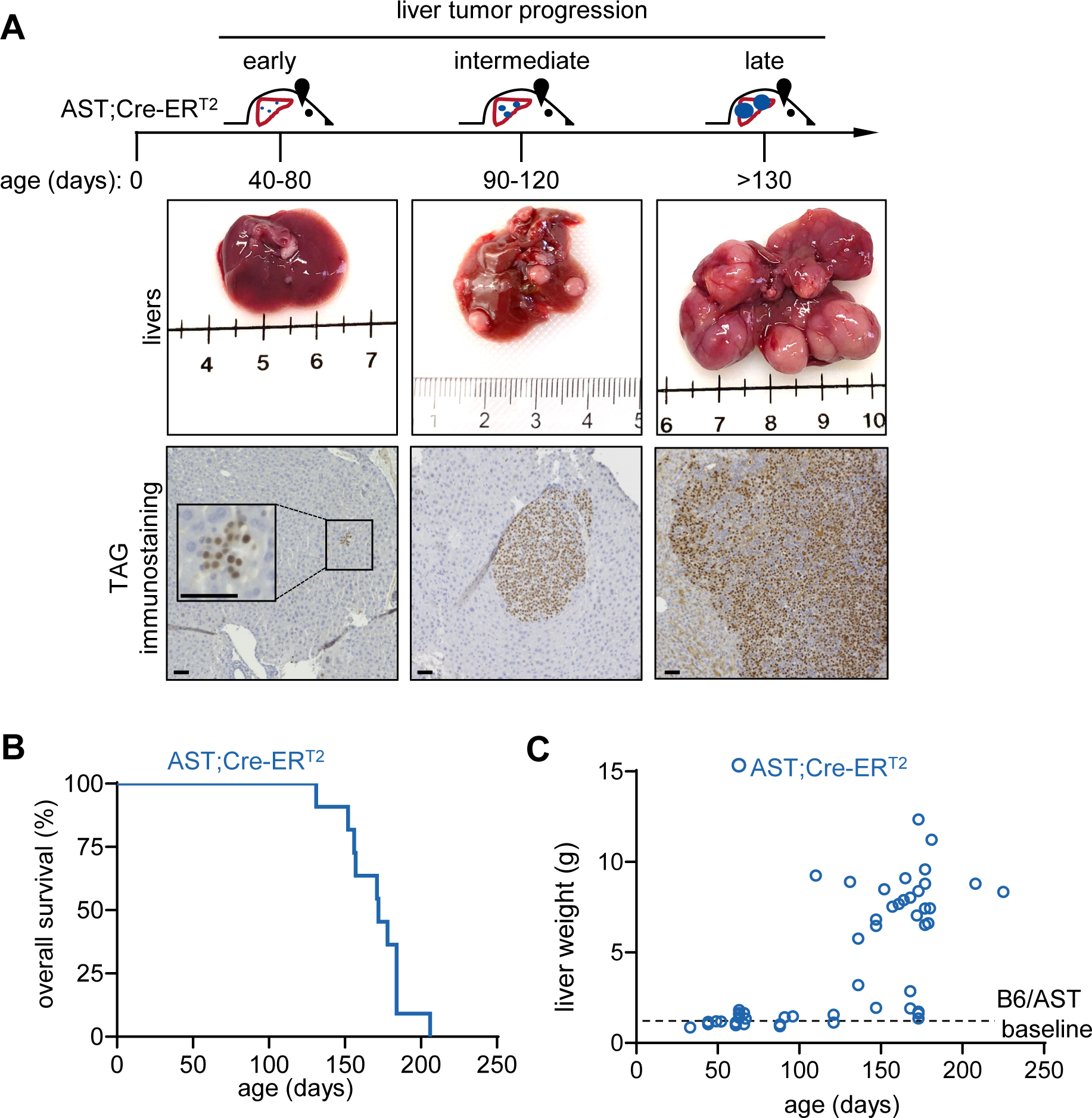
AST;Cre-ER^T2^ mice sporadically develop liver tumors which progress with age. **A.** Timeline of tumor development in AST;Cre-ER^T2^ mice, with the middle panel showing corresponding gross liver images (ruler measured in centimeters (cm)) and the lower panel showing TAG immunohistochemistry staining (scale bar = 50 μ). AST;Cre-ER^T2^ mice aged 40-80d with grossly normal liver and few TAG+ hepatocytes are classed as “early”; age 90-120d mice with small macroscopic liver tumors and larger foci of TAG+ hepatocytes are classed as “intermediate”; >130d mice with large tumors occupying most of the liver and large TAG+ lesions are “late.” Representative gross and TAG immunohistochemistry images of early, intermediate, and late AST;Cre-ER^T2^ mice are shown. **B.** Kaplan-Meier curve showing overall survival of AST;Cre-ER^T2^ mice. **C.** Liver weights of AST;Cre-ER^T2^ male (n=59) mice plotted against age. Black dotted line shows baseline weights of tumor-free B6 and AST mice.

### TST activate and proliferate similarly in mice with early and late lesions

To compare initial TST differentiation in mice with early versus late liver lesions, we transferred carboxy fluorescein succinimidyl ester (CFSE)-labeled naive TCR_TAG_ into early and late time point AST;Cre-ER^T2^ mice (**Fig. 2A**). TCR_TAG_ underwent robust proliferation in mice with early or late liver lesions and upregulated PD1 as they divided (**Fig. 2B, lower panels**). TCR_TAG_ in mice with early lesions lagged 1-2 divisions behind TCR_TAG_ in the spleen, liver-draining lymph node (ldLN), and livers of mice with late lesions (**Fig. 2B, C**), and we found fewer TCR_TAG_ in the spleens, ldLN, and livers of early mice as compared to late mice (**Fig. 2D**), likely reflecting the lower antigen burden in early versus late mice. Strikingly, within 60h of transfer, most TCR_TAG_ in both early and late mice failed to produce effector cytokines TNFα and IFNγ (**Fig. 2E)**. However, in mice with early liver lesions we identified a small population of TCR_TAG_ in the spleen and liver that could produce effector cytokines (**Fig. 2E**). Thus, in hosts with sporadic early lesions, a small but signficant subset of TST resisted rapid differentiation to the dysfunctional state.

**Figure 2:**
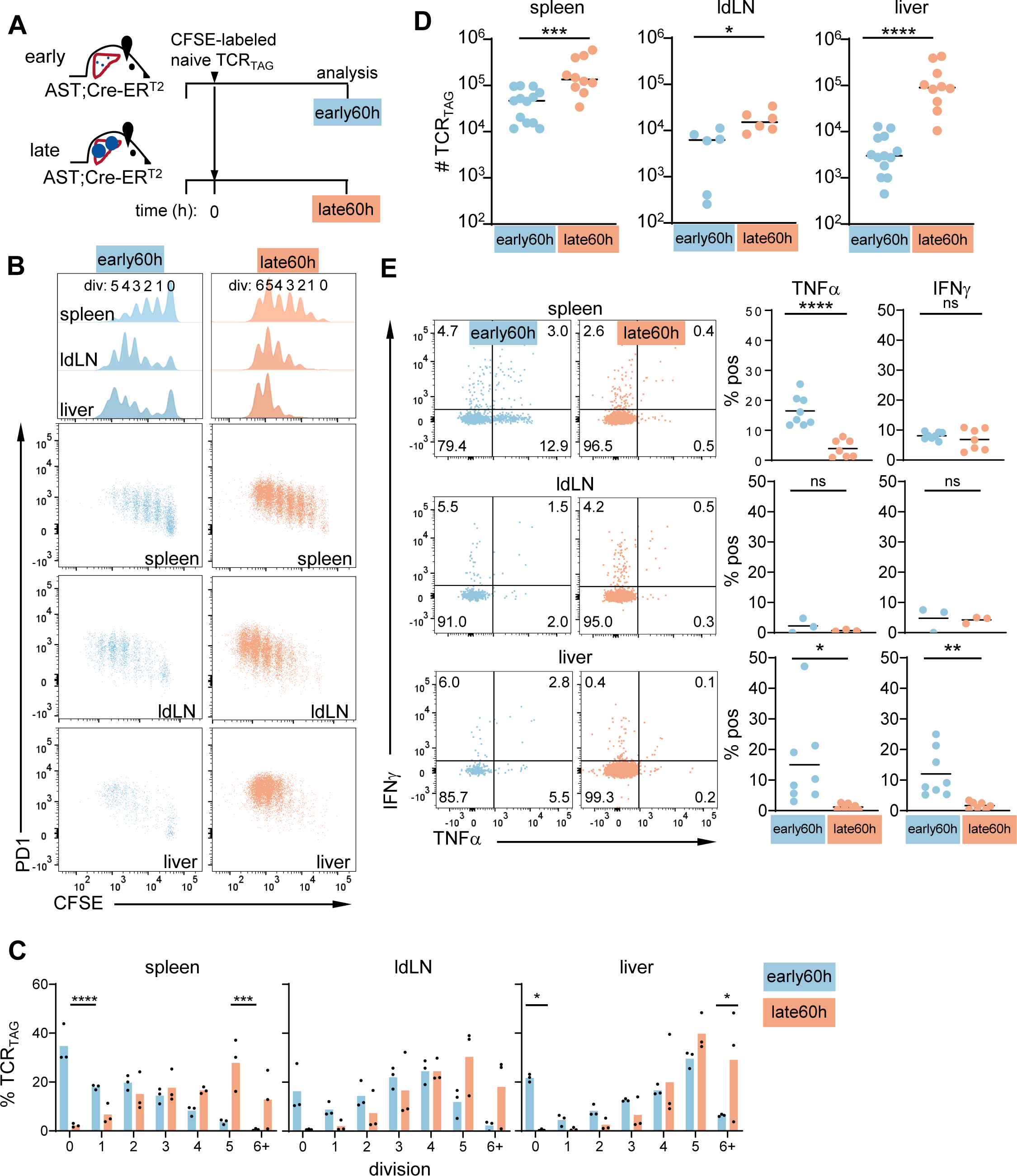
TST activate and proliferate similarly in mice with early and late lesions. **A.** Scheme: CFSE-labeled TCR_TAG_ cells were adoptively transferred into early (blue) and late (orange) AST;Cre-ER^T2^ mice and analyzed 60 hours (h) later. **B.** TCR_TAG_ CFSE dilution and PD1 versus CFSE dilution in the spleens, liver-draining lymph nodes (ldLN) and livers of early and late mice. This and all subsequent flow plots are gated on live CD8+ Thy1.1+ cells (representative gating is shown in **Supplemental** Fig. 1A). Data is concatenated from 3 biologic replicates and representative of three independent experiments. **C**. Percentage of TCR_TAG_ cells by cell division for spleens, ldLNs, and livers. Each symbol represents an individual mouse with n=3/group. \**P*<0.05, \*\*\**P*<0.001, \*\*\*\**P*<0.0001 (two-way ANOVA, followed by post-hoc Šidák test.) Data is representative of two independent experiments. **D.** Number of TCR_TAG_ in spleens, ldLN, and livers of early and late mice. Each symbol represents an individual mouse with n=10-13/group combined from three independent experiments for spleens and livers, and n=6 combined from two independent experiments for ldLN. \**P*<0.05, \*\*\**P*<0.001, \*\*\*\**P*<0.0001 (unpaired Student t test). **E.** Left, TCR_TAG_ TNFα and IFNγ production after 4-hour *ex vivo* TAG peptide stimulation, assessed by flow cytometry. Data is concatenated from three biologic replicates and representative of two independent experiments for spleens and livers, and one independent experiment for ldLN. Right, percentage of TCR_TAG_ positive for TNFα and IFNγ. Gates for this and subsequent cytokine production figures are set based on “no peptide stimulation” control. Each symbol represents an individual mouse with n=7-8/group for spleen and liver and n=3/group for ldLN. ns=not significant, \**P*<0.05, \*\**P*<0.01, \*\*\*\**P*<0.0001 (unpaired Student t test).

### A subset of TST remain functional in mice with early liver lesions

To determine whether this functional TST subset persisted, we examined TCR_TAG_ immunophenotype and function 5d and 21d post transfer into early or late AST;Cre-ER^T2^ mice (**Fig. 3A**). Though late lesion mice initially had higher numbers of TCR_TAG_, by 21d TCR_TAG_ numbers were similar in early and late lesion mice (**Fig. 3B**). TCR_TAG_ in both early and late mice upregulated CD44, indicating antigen exposure and activation (**Supplemental Fig. 2**). Notably, TCR_TAG_ in early mice upregulated PD1 expression, suggesting that PD1 expression can identify tumor-reactive TST even in hosts with early lesions (**Fig. 3C, upper panels**). PD1+ TST which express the transcription factor (TF) TCF1 and are localized to secondary lymphoid organs or tertiary lymphoid structures have been associated with improved anti-tumor function and responsiveness to immunotherapy^9^ ^17^ (reviewed^3^ ^11^). At 21d, TCR_TAG_ in early mice expressed higher levels of TCF1 in the spleen (**Fig. 3C**, **lower panels**). While nearly all TCR_TAG_ in mice with late lesions were dysfunctional and failed to produce effector cytokines, mice with early lesions retained a subset of IFNγ-producing TCR_TAG_, even at 21d (**Fig. 3D**). These findings are in line with previous work in a mouse model of sporadic cancer arising in all organs which found that early immune tolerance was responsible for tumor progression^18^.

**Figure 3:**
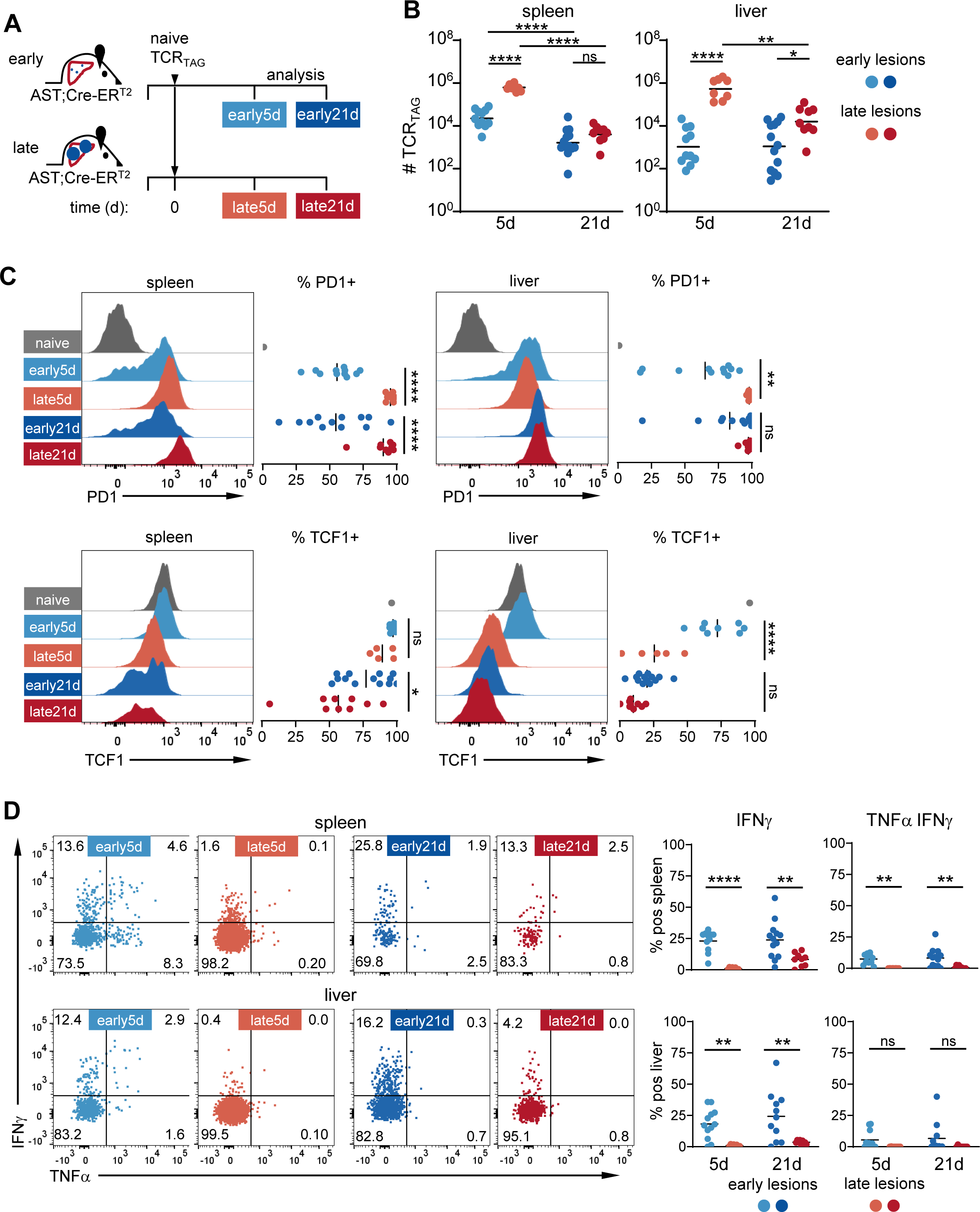
A subset of TST remain functional in mice with early liver lesions. **A.** Scheme: TCR_TAG_ Thy1.1 (21day (d)) and Thy1.12 cells (5d) were sequentially adoptively transferred into early (blue) and late (red) AST;Cre-ER^T2^ mice and analyzed simultaneously post-transfer. **B.** Number of TCR_TAG_ in spleens and livers of early and late mice. \**P*<0.05, \*\**P*<0.01, \*\*\*\**P*<0.0001 (two-way ANOVA with post-hoc Tukey test). **C.** Left, histograms of TCR_TAG_ PD1 and TCF1 expression. Right, percentage of PD1+ and TCF1+ TCR_TAG_ with positive gate set to exclude (PD1) or include (TCF1) naive TCR_TAG_ (gray) (representative gating scheme is shown in **Supplemental** Fig. 1B). \**P*<0.05, \*\**P*<0.01, \*\*\**P*<0.001 (two-way ANOVA followed by post-hoc Šídák test). **D.** Left, TCR_TAG_ TNFα and IFNγ production after 4-hour *ex vivo* TAG peptide stimulation. Right, percentage of TCR_TAG_ positive for IFNγ and TNFα/IFNγ. \*\**P*<0.01, \*\*\*\**P*<0.0001 (two-way ANOVA followed by post-hoc Šídák test). For B-D, each symbol represents an individual mouse with n=11-13/early group and n=8-9/late group and combined from three independent experiments.

### Vaccination early during tumorigenesis halts cancer progression

We next asked if the functional TST subset present in mice with early lesions could be harnessed to stop tumor progression. *Listeria monocytogenes* (LM) is a gram-positive intracellular bacterium which induces strong CD4 and CD8 T cell responses^19^. Using an *actA inlB* deficient attenuated LM vaccination strain^20^, which has been used in human clinical trials for advanced cancers^21–23^, and engineered to express the TAG epitope I (LM_TAG_)^5^, we tested whether early vaccination of AST;Cre-ER^T2^ would protect mice from liver cancer progression. Young AST;Cre-ER^T2^ mice adoptively-transferred with naive TCR_TAG_ were either untreated (Φ), given a single dose of empty LM (LMΦ), or vaccinated with a single dose of LM_TAG_ and followed, with some cohorts analyzed prior to endpoint (**Fig. 4A**). LM_TAG_-immunization conferred a major survival advantage, with all mice remaining tumor-free and one mouse euthanized for dermatitis without any evidence of liver tumors (**Fig. 4B, C** and **Supplemental Fig. 3A**). In contrast, most mice in the Φ and LMΦ groups reached endpoint with multiple large liver tumors and increased liver weight (**Fig. 4B, C**). At endpoint, TCR_TAG_ in the LM_TAG_-immunized mouse made effector cytokines, in contrast to the TCR_TAG_ in the tumor-bearing mice in the Φ and LMΦ groups which did not (**Supplemental Fig. 3B**), suggesting that functional TCR_TAG_ prevent liver tumor progression.

**Figure 4:**
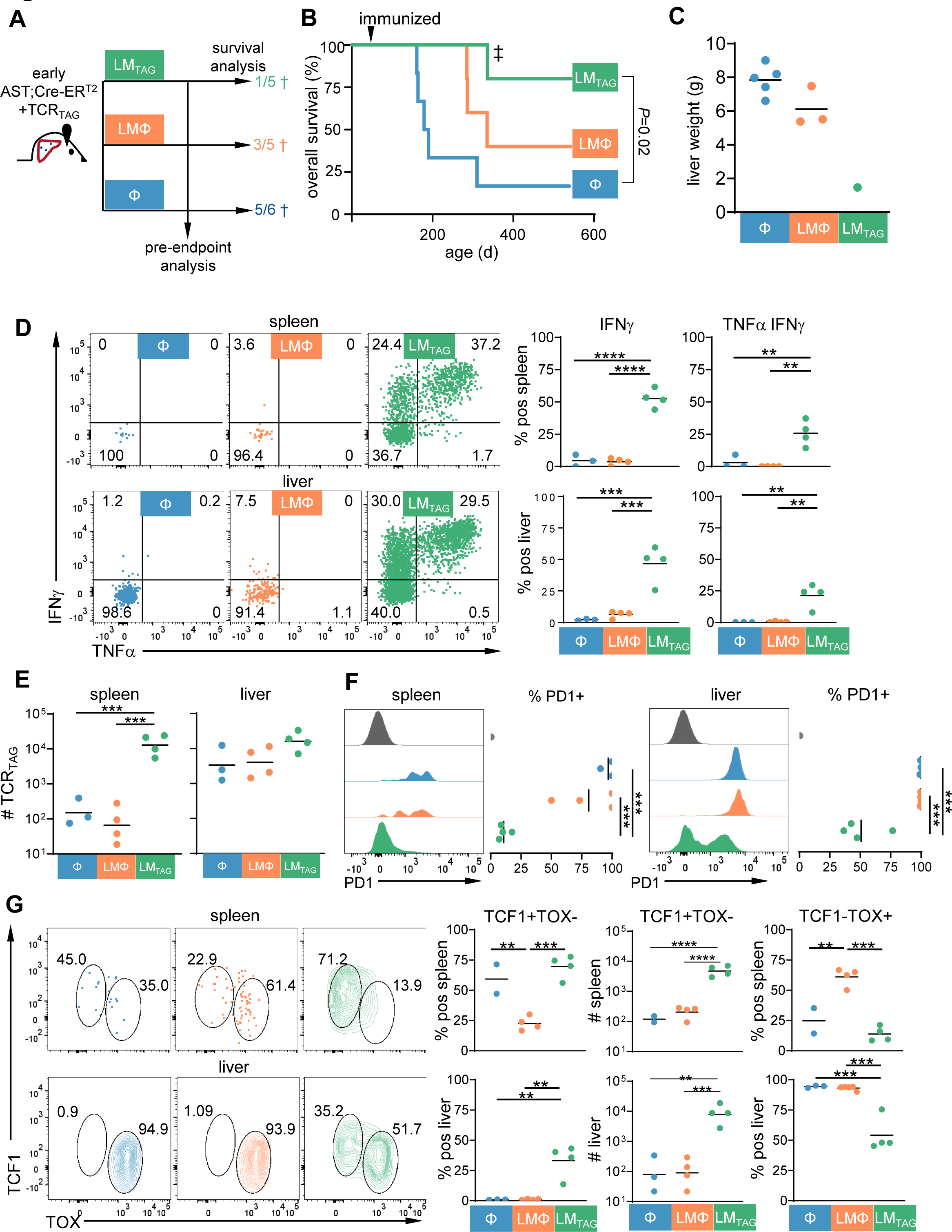
Vaccination early during tumorigenesis halts cancer progression. **A**. Scheme: Early (age 42d) AST;Cre-ER^T2^ mice were adoptively transferred with TCR_TAG_ and untreated (Φ), treated with empty LM (LMΦ), or vaccinated with LM expressing the TAG antigen (LM_TAG_), with one cohort followed to endpoint and the second analyzed at age 100d prior toendpoint. The numbers indicate the number of mice in the long-term cohort who reached endpoint. **B.** Kaplan-Meier curve showing survival of mice in each group with n=5-6/treatment group. Statistical analysis was performed using log rank (Mantel-Cox). The ‡ indicates a mouse euthanized for dermatitis, whereas all other mice were euthanized due to advanced liver tumors. **C.** Liver weights of euthanized mice at end point. Each symbol represents an individual mouse with n=1-5/group. **D-G.** Cohort as in A was analyzed at age 100d prior to endpoint. **D**. Left, TCR_TAG_ TNFα and IFNγ production after 4-hour *ex vivo* TAG peptide stimulation. Right, percentage of TCR_TAG_ positive for IFNγ and TNFα/IFNγ. Each symbol represents an individual mouse with n=3-4/group. \**P*<0.05 (one-way ANOVA with post-hoc Tukey test). **E.** Number of TCR_TAG_ in spleen and livers. Each symbol represents an individual mouse with n=3-4/group. \*\*\**P*<0.001 (one-way ANOVA with post-hoc Tukey test). **F**. Left, histograms of TCR_TAG_ PD1 expression. Right, percentage of PD1+ TCR_TAG_ with positive gate set to exclude naive TCR_TAG_ (gray). Each symbol represents an individual mouse with n=3-4/group. \*\*\**P*<0.001 (one-way ANOVA with post-hoc Tukey test). **G.** Left, TCF1 and TOX expression in TCR_TAG_ cells. Data is concatenated from 2-4 biologic replicates/group. Right, percentage of TCF1+TOX- and TCF1-TOX+ TCR_TAG_. Each symbol represents an individual mouse with n=2-4/group. \**P*<0.05, \*\*\**P*<0.001 (one-way ANOVA with post-hoc Tukey test). Data in D-G is representative of two independent experiments.

### Progenitor TST are associated with vaccine efficacy

To determine whether the presence of functional TST correlated with later tumor-free survival, we analyzed TCR_TAG_ in Φ, LMΦ, or LM_TAG_-vaccinated mice at age 100d, before mice reached endpoint. TCR_TAG_ in the spleen and livers of Φ or LMΦ-treated mice made no effector cytokine (**Fig. 4D**), suggesting that without vaccination, the small IFNγ-producing TST subset observed at the earlier 21d time point failed to persist. In stark contrast, mice in the LM_TAG_ group had a large subset of double-producing TNFα^+^IFNγ^+^ TCR_TAG_ in both the spleen and liver (**Fig. 4D**). LM_TAG_-immunized mice had a larger population of TCR_TAG_ in the spleen (**Fig. 4E**), which were PD1**^-^** (**Fig. 4F**, left panel). Interestingly, there were two population of TCR_TAG_ in the livers of LM_TAG_-vaccinated mice, a PD1^-^ and a PD1^int^ population (**Fig. 4F**, right panel). We assessed expression of TCF1 and TOX, a TF associated with terminally-differentiated TST^24–28^. LM_TAG_-vaccinated mice had a higher number of TCF1+TOX-progenitor TST in the spleen and liver as compared to Φ and LMΦ-treated mice (**Fig. 4G**). While most TST in LM_TAG_-vaccinated spleens were TCF1+TOX-, in the liver there were two populations, a TCF1+TOX- and a TCF1-TOX+ population (**Fig. 4G**, lower panel). Taken together, these data suggest that LM_TAG_ vaccination induces a robust population of progenitor TST, which sustains long-term anti-tumor responses, eliminating transformed hepatocytes and blocking cancer progression.

We next tested whether later vaccination could confer similar protection against liver tumor progression (**Supplemental Fig. 3C**). Immunization at an intermediate time point (100d), when few progenitor TST exist (Φ group, Fig. 4G), failed to slow liver tumor progression, and mice succumbed to tumors between 150-175d (**Supplemental Fig. 3D, E**), suggesting that progenitor TST are required for vaccine anti-tumor efficacy.

### Vaccination is superior to ICB in blocking tumor progression

An important and open question in cancer immunotherapy is how ICB versus vaccination compares in reprogramming anti-cancer immune responses and how best to combine and sequence these therapies^2^ ^29^. Therefore, we compared the efficacy of ICB, LM_TAG_ vaccination, and combined ICB/ LM_TAG_ vaccination. Early AST;Cre-ER^T^^2^ mice adoptively transferred with TCR_TAG_ were treated with isotype control antibodies (iso), anti-PD1/anti-PD-L1 antibodies (ICB), LM_TAG_, or combined LM_TAG_/ICB (**Fig. 5A**). ICB showed no benefit as compared to iso, with all mice developing large liver tumors (**Fig. 5B, C**). In contrast, LM_TAG_ and LM_TAG_/ICB-treated mice had no evidence of tumor progression at >350d (**Fig. 5B**). We next analyzed TCR_TAG_ in iso, ICB, LM_TAG_, or ICB/LM_TAG_-treated mice at age 100d. LM_TAG_ vaccination, either alone or in combination, led to a substantial increase in TST numbers and IFNγ production, while ICB alone had little impact (**Fig. 5D, E and Supplemental Fig. 4A, B**). Notably, the addition of ICB to LM_TAG_ did not improve TCR_TAG_ number or function (**Fig. 5D, E and Supplemental 4A, B**).

**Figure 5:**
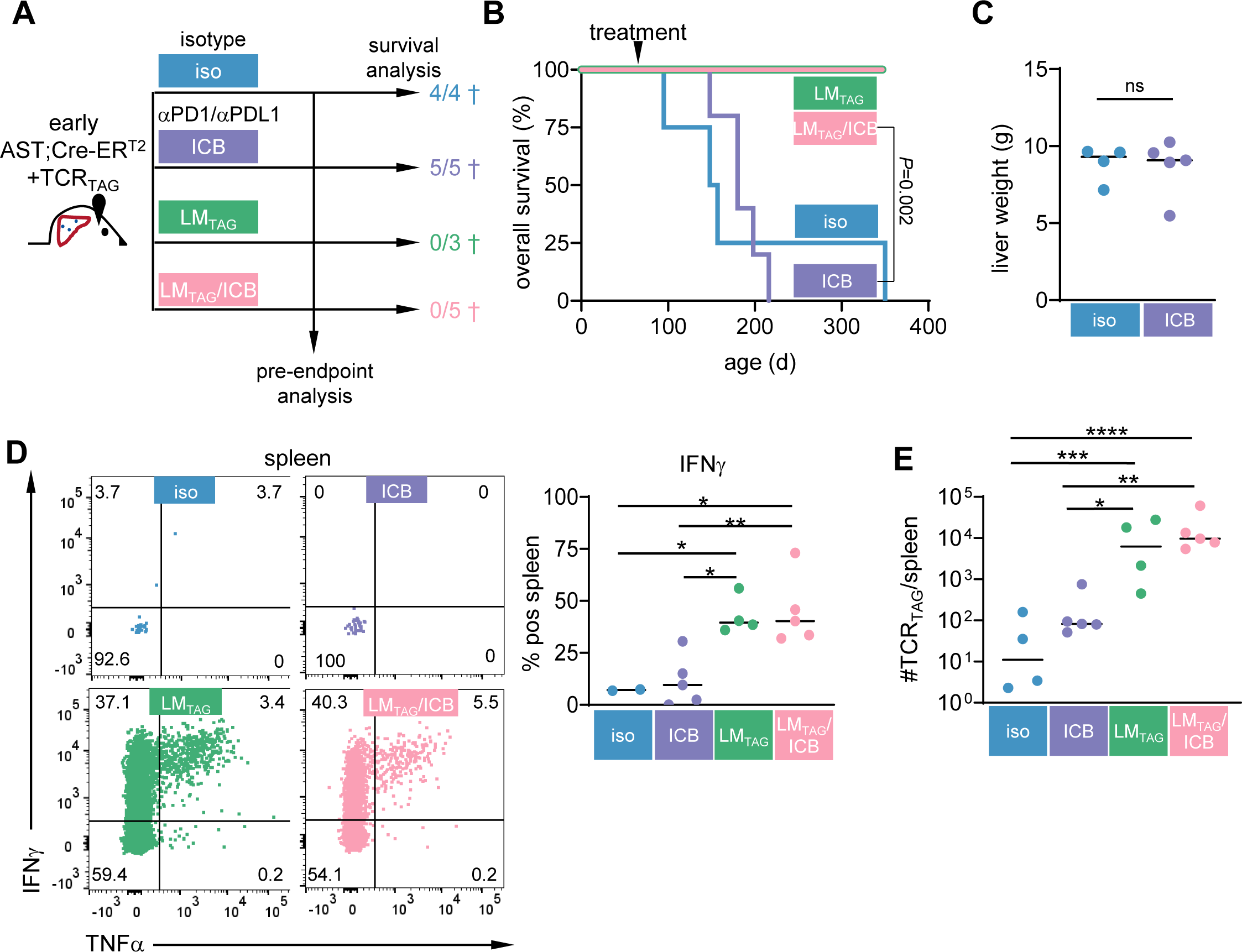
Vaccination is superior to ICB in blocking tumor progression. **A**. Scheme: Early AST;Cre-ER^T2^ mice were adoptively transferred with TCR_TAG_ and treated with isotype control antibody, anti-PD1/PDL1 antibodies (ICB), LM_TAG_, or combination LM_TAG_/ICB, with one cohort followed to endpoint and the second analyzed at age 100d prior to endpoint. The numbers indicate the number of mice in the long-term cohort who reached endpoint. **B**. Kaplan-Meier curve showing survival of mice in each group. Statistical analysis was performed using log rank (Mantel-Cox) test with Bonferroni correction for multiple comparisons. **C.** Graph showing liver weights at endpoint. Each symbol represents an individual mouse with n=4-5/group (unpaired Student t test). **D-E.** Cohort as in A was analyzed at age 100d prior to endpoint. **D**. Left, spleen TCR_TAG_ TNFα and IFNγ production after 4-hour *ex vivo* TAG peptide stimulation. Right, percentage of TCR_TAG_ positive for IFNγ. Each symbol represents an individual mouse with n=2-5/group. \**P*<0.05, \*\**P*<0.01 (one-way ANOVA followed by post-hoc Tukey test). **E.** Number of TCR_TAG_ cells in the spleen. \**P*<0.05, \*\**P*<0.01, \*\*\**P*<0.001, \*\*\*\**P*<0.0001 (one-way ANOVA followed by post-hoc Tukey test).

## DISCUSSION

Here, we utilized an autochthonous liver cancer mouse model in which hepatocytes undergo spontaneous oncogene activation at a low frequency. With time, transformed hepatocytes progress from microscopic to small macroscopic to large late-stage liver tumors. Interestingly, while most TST activated in mice with early lesions upregulated PD1 and lost the ability to produce effector cytokines, a small subset of PD1^+^TCF1^+^ TST persisted and retained the ability to produce IFNγ. Without intervention, this functional subset failed to persist, and the mice developed tumors. However, vaccination of early mice generated a large population of progenitor and polyfunctional TST, which blocked tumor progression and resulted in long-term survival. Given that the TAG oncogene is encoded in the germline, putting every hepatocyte at risk of transformation, this long-term protection is striking. In contrast, ICB treatment failed to confer protection from liver cancer progression or augment LM_TAG_ vaccination. Of note, later vaccination, at a time when the progenitor population was no longer present, failed to generate a polyfunctional population or block tumor progression.

The development and use of cancer vaccines is a key strategy to prevent or halt cancer progression and a main goal of the National Cancer Plan (https://nationalcancerplan.cancer.gov/about). LM-based vaccines have been tested in clinical trials with poor or mixed results^19^. These studies have mainly tested LM vaccines in patients with advanced or refractory cancers. Our studies could provide mechanistic insigh as to why vaccine in patients with advanced cancers fail: for vaccines to be effective, a progenitor TST progenitor population must be present. We previously showed that over time and with continued tumor antigen exposure, the TCF1+ TST population diminishes^17^, and there is increasing evidence that tumor burden negatively correlates with responses to ICB (reviewed in ^30^) and chimeric antigen receptor T cell therapy^31^. Hosts with early lesions and lower tumor antigen burden are more likely to harbor a population of progenitor TCF1+ TST that respond to vaccination and protect against further tumor progression.

Our finding that vaccination blocked tumor progression while ICB did not may be surprising at first-glance, given that ICB is also known to act by recruiting progenitor TST to mediate anti-tumor responses^32^ ^33^. An important point demonstrated by our results (Fig. 3) and previous studies is that not all TCF+ TST are functional, nor does ICB alone lead to functional reprogramming^17^ ^34^. TST activated in tumor-bearing hosts rapidly undergo widespread epigenetic remodeling, which is reinforced with continued tumor antigen exposure^16^. Removal from tumor^16^ or PD-1 blockade^35^ is not sufficient to reverse dysfunctional epigenetic reprogramming. Our findings suggest that LM_TAG_ vaccination, which provides antigenic stimulation in the context of an immunogenic pathogen, can reprogram dysfunctional progenitor TCF1^+^ TST. Future studies will be needed to decipher mechanisms by which LM_TAG_ vaccination reprogram progenitor TST, why terminally-differentiated TST fail to respond, and how different immunotherapies such as vaccines and ICB can be optimally combined. The sporadic liver cancer model we have developed and characterized provides an excellent platform for such future investigations, which could enable us to make long-term progression-free survival a reality for all patients with cancer.

## DECLARATIONS

### Ethics approval

All experiments were approved by the Vanderbilt University Medical Center (VUMC) Institutional Animal Care and Use Committee.

### Patient consent for publication

Not applicable.

### Availability of data and material

All data relevant to the study are included in the article or uploaded as online supplemental information. The data used and/or analyzed during the current study are available from the corresponding author on reasonable request.

## Author’s contributions

Conception and design: CRDR and MP. Experiments: CRDR, MWR, FR, NRF, KAM, JJR, MME. Analysis and interpretation of data: CRDR and MP. Writing of the manuscript: all authors.

## Acknowledgements

We thank the Vanderbilt Division of Animal Care, the Vanderbilt Translational Pathology Core, and the VUMC DDRC Tissue Morphology Core. We thank Dr. Peter Lauer and Aduro Biotech (since acquired by Chinook Therapeutics) for providing attenuated *Listeria* strains.

## Authors’ information

ORCID Mary Philip https://orcid.org/0000-0001-7496-2630 Twitter@Philiplabvandy

---

**Supplemental Figure 1: Representative gating strategy flow analysis. A.** Representative gating scheme to identify and analyze Thy1.1 and/or Thy1.12 TCR_TAG_ cells isolated from livers (shown) and spleen. **B.** Representative gating scheme to determine the percentage of PD1+ and TCF1+ TCR_TAG_. The positive gate was set to exclude naive (PD1, left) or include naive (TCF1, right) TCR_TAG_ (gray).

**Supplemental Figure 2: TCR_TAG_ upregulate CD44 in mice with early and late lesions.** Left, histograms of TCR_TAG_ CD44 expression. Right, percentage of CD44+ TCR_TAG_ with positive gate set to exclude naive TCR_TAG_ (gray). Each symbol represents an individual mouse with n=11-13/early and n=8-9/late combined from three independent experiments. ns=not significant, \**P*<0.05, \*\*\*\**P*<0.0001 (two-way ANOVA with post-hoc Tukey test).

**Supplemental Figure 3: Early but not later vaccination prevents liver tumor progression. A.** Gross liver image from LM_TAG_-vaccinated mouse euthanized for dermatitis at 336d (ruler measured in cm). **B.** Left, TCR_TAG_ TNFα and IFNγ production after 4-hour *ex vivo* TAG peptide stimulation at endpoint. Right, percentage of TCR_TAG_ positive for IFNγ and TNFα/IFNγ. Each symbol represents an individual mouse with n=4-5/group. C. Early AST;Cre-ER^T2^ mice were adoptively transferred with TCR_TAG_ and immunized at age 100d with LMΦ or LM_TAG_ and followed. The numbers indicate the number of mice who reached endpoint. D. Kaplan-Meier curve showing survival of mice in each group. Statistical analysis was performed using log rank (Mantel-Cox) test. E. Graph showing liver weights at endpoint. Each symbol represents an individual mouse with n=4/group. Statistical analysis was performed using unpaired Student t test.

**Supplemental Figure 4: Vaccination is superior to ICB in blocking tumor progression.** Mice were treated as in Fig. 5A and analyzed at age 100d prior to endpoint. **A.** Left, liver TCR_TAG_ TNFα and IFNγ production after 4-hour *ex vivo* TAG peptide stimulation. Right, percentage of TCR_TAG_ positive for IFNγ. Each symbol represents an individual mouse with n=4-5. \**P*<0.05 (one-way ANOVA followed by post-hoc Tukey test). **B.** Number of TCR_TAG_ cells in the liver. \**P*<0.05, \*\*\**P*<0.001 (one-way ANOVA followed by post-hoc Tukey test).

