## Supplemental Figures for "Vaccination generates functional progenitor tumor-specific CD8 T cells and long-term tumor control"

### Supplemental Figure 1

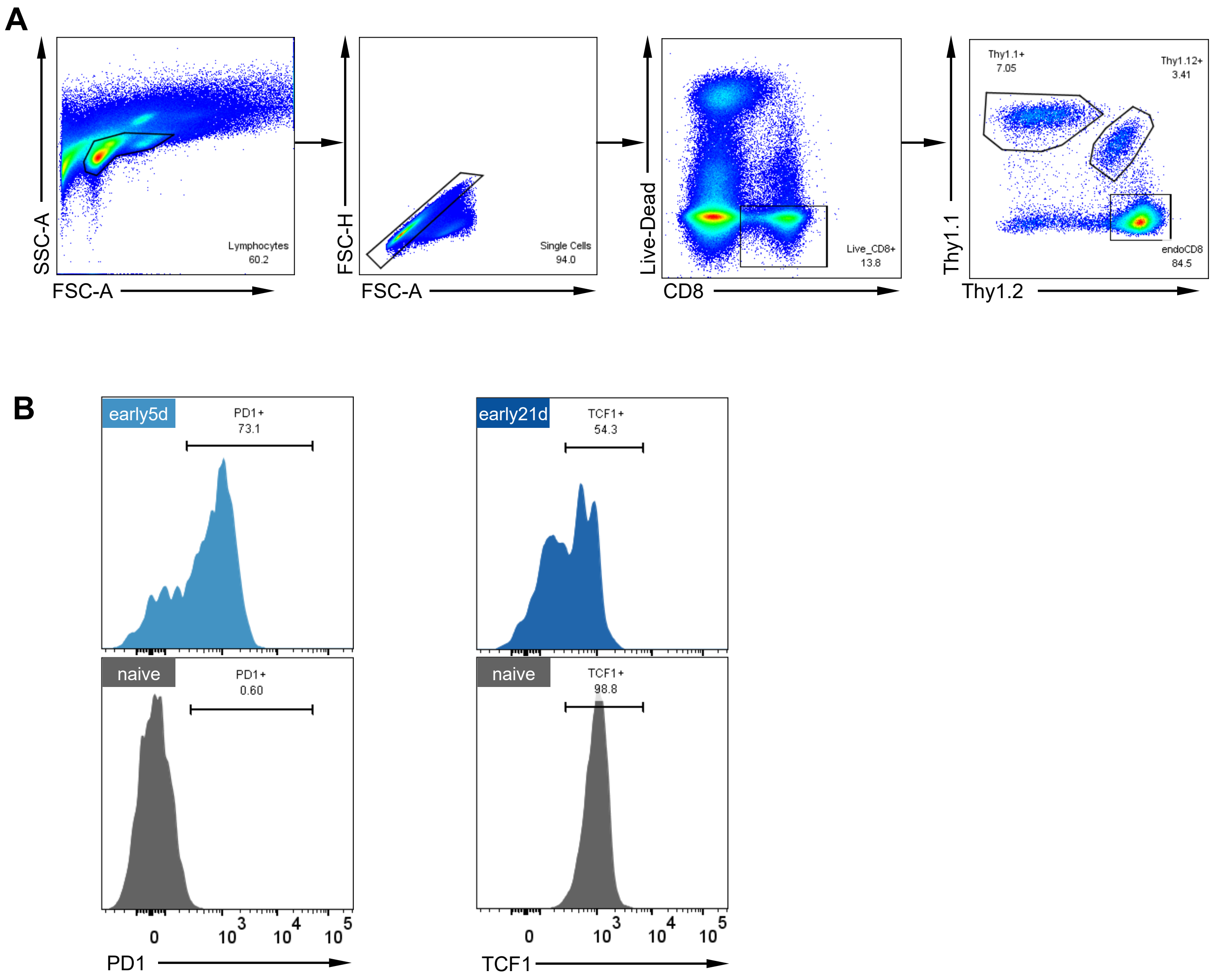


**Supplemental Figure 1: Representative gating strategy flow analysis. A.** Representative gating scheme to identify and analyze Thy1.1 and/or Thy1.12 $TCR_{TAG}$ cells isolated from livers (shown) and spleen. **B.** Representative gating scheme to determine the percentage of PD1+ and TCF1+ $TCR_{TAG}$. The positive gate was set to exclude naive (PD1, left) or include naive (TCF1, right) $TCR_{TAG}$ (gray).

### Supplemental Figure 2

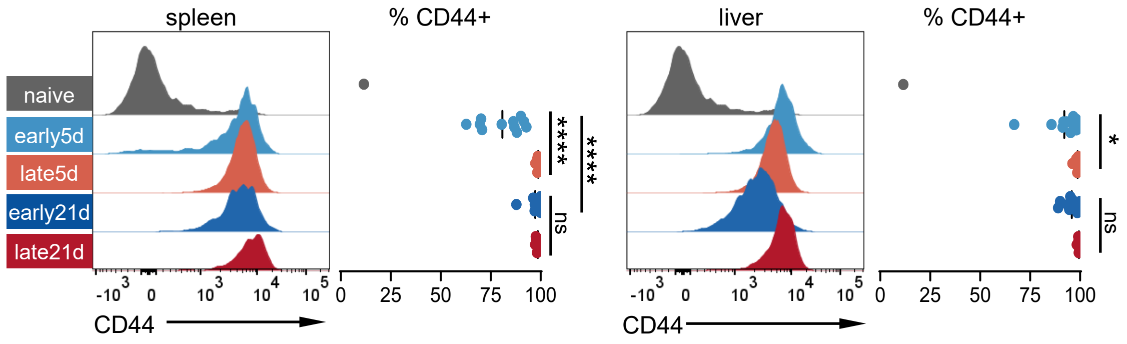


**Supplemental Figure 2: $TCR_{TAG}$ upregulate CD44 in mice with early and late lesions.** Left, histograms of $TCR_{TAG}$ CD44 expression. Right, percentage of CD44+ $TCR_{TAG}$ with positive gate set to exclude naive $TCR_{TAG}$ (gray). Each symbol represents an individual mouse with n=11-13/early and n=8-9/late combined from three independent experiments. ns=not significant, **P*<0.05, *****P*<0.0001 (two-way ANOVA with post-hoc Tukey test).

### Supplemental Figure 3

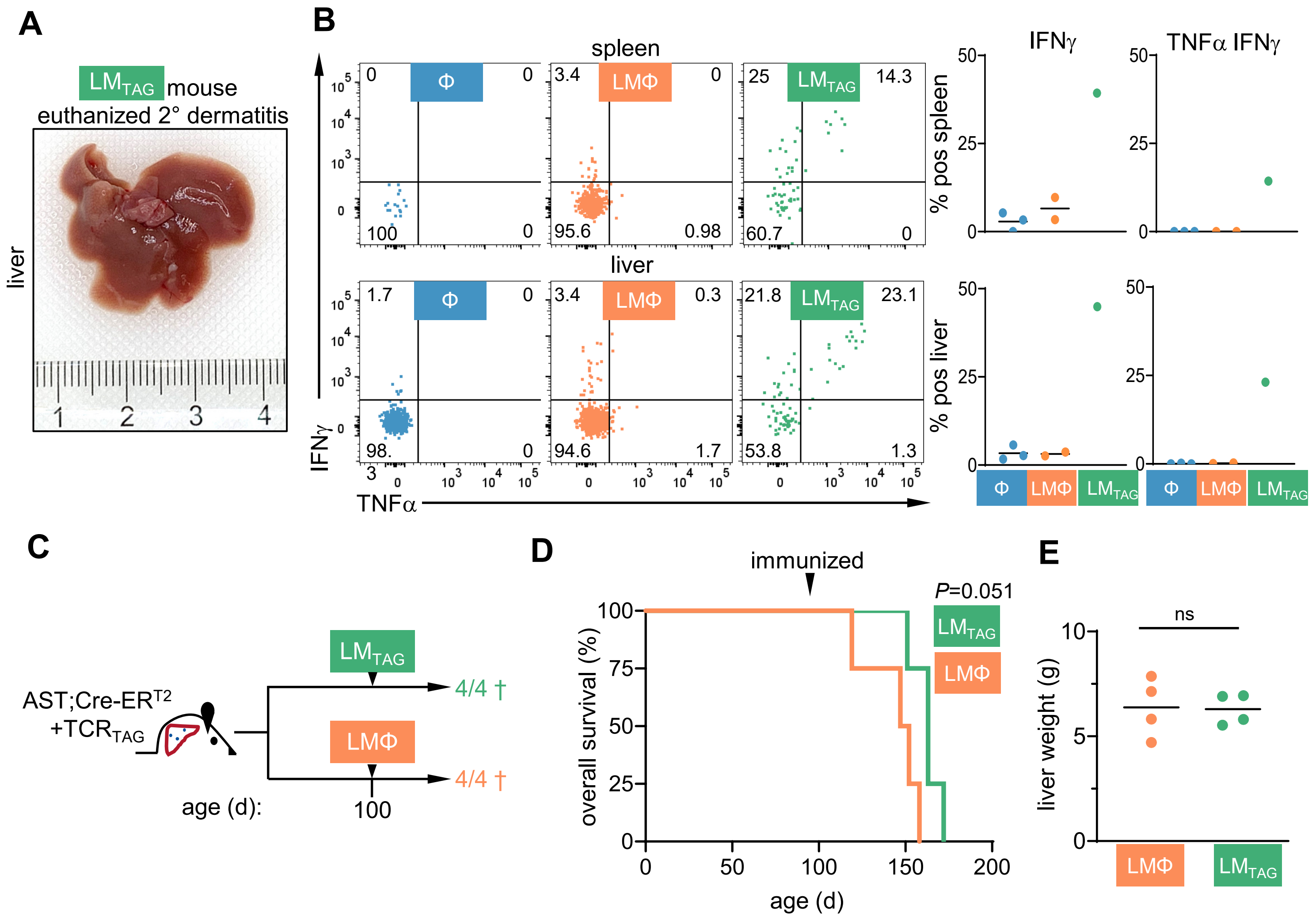


**Supplemental Figure 3: Early but not later vaccination prevents liver tumor progression. A.** Gross liver image from $LM_{TAG}$-vaccinated mouse euthanized for dermatitis at 336d (ruler measured in cm). **B.** Left, $TCR_{TAG}$ TNFα and IFNγ production after 4-hour *ex vivo* TAG peptide stimulation at endpoint. Right, percentage of $TCR_{TAG}$ positive for IFNγ and TNFα/IFNγ. Each symbol represents an individual mouse with n=4-5/group. **C.** Early AST;Cre-$ER^{T2}$ mice were adoptively transferred with $TCR_{TAG}$ and immunized at age 100d with LMΦ or $LM_{TAG}$ and followed. The numbers indicate the number of mice who reached endpoint. **D.** Kaplan-Meier curve showing survival of mice in each group. Statistical analysis was performed using log rank (Mantel-Cox) test. **E.** Graph showing liver weights at endpoint. Each symbol represents an individual mouse with n=4/group. Statistical analysis was performed using unpaired Student t test.

### Supplemental Figure 4

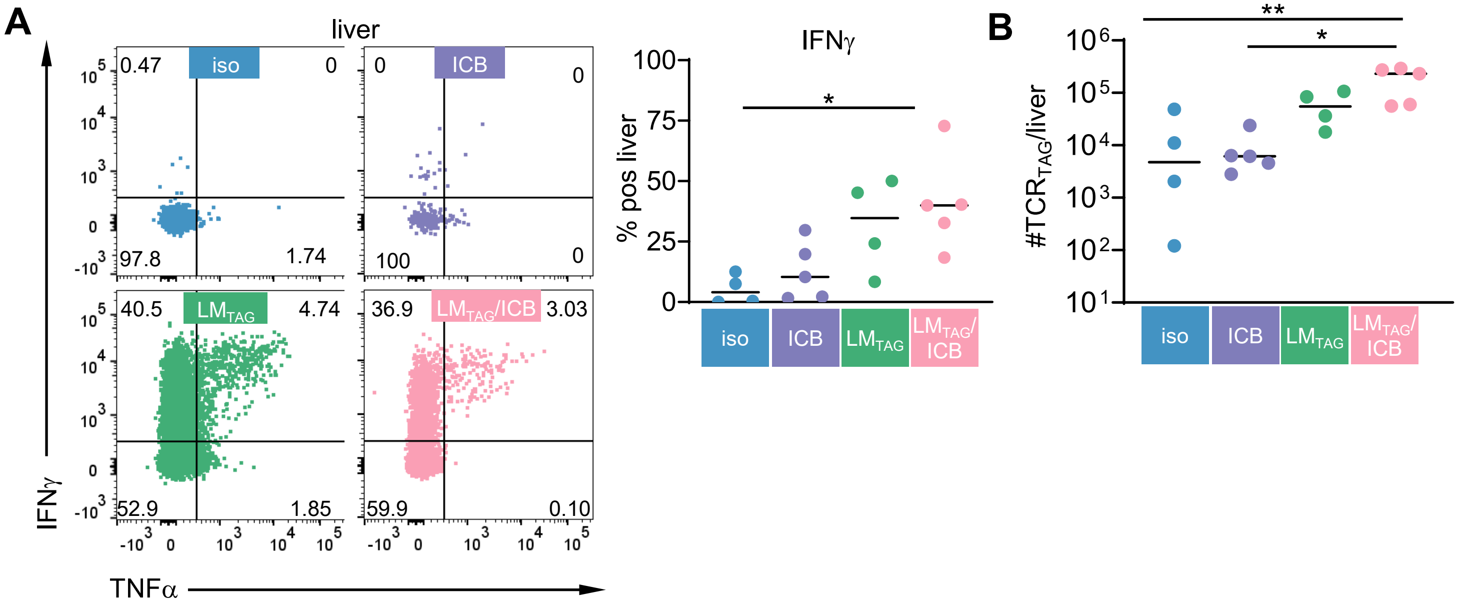


**Supplemental Figure 4: Vaccination is superior to ICB in blocking tumor progression.** Mice were treated as in Fig. 5A and analyzed at age 100d prior to endpoint. **A.** Left, liver $TCR_{TAG}$ TNFα and IFNγ production after 4-hour *ex vivo* TAG peptide stimulation. Right, percentage of $TCR_{TAG}$ positive for IFNγ. Each symbol represents an individual mouse with n=4-5. *$P$<0.05 (one-way ANOVA followed by post-hoc Tukey test). **B.** Number of $TCR_{TAG}$ cells in the liver. *$P$<0.05, ***$P$<0.001 (one-way ANOVA followed by post-hoc Tukey test).
