## Supplemental Tables for "Vaccination generates functional progenitor tumor-specific CD8 T cells and long-term tumor control"

**Table 1.** Flow cytometry antibodies.

| Antibody | Fluorophore | Clone | Source | Identifier |
|---|---|---|---|---|
| *Anti-CD8a* | BV605 | 53-6.7 | BioLegend | Cat# 100744 |
| *Anti-CD44* | PcP-Cy5.5 | IM7 | Tonbo | Cat# 65-0441 |
| *Anti-CD44* | FITC | IM7 | BioLegend | Cat# 103006 |
| *Anti-Thy1.1* | BV510 | OX-7 | BioLegend | Cat# 202535 |
| *Anti-Thy1.2* | BV421 | 53-2.1 | BioLegend | Cat# 140327 |
| *Anti-IFNγ* | APC | XMG.12 | BioLegend | Cat# 505810 |
| *Anti-PD1* | APC | RMP1-30 | BioLegend | Cat# 109112 |
| *Anti-TCF1* | AF647 | C63D9 | Cell Signaling Technology | Cat# 6709S |
| *Anti-TNFα* | PE | MP6-XT22 | Life | Cat# 12-7321-82 |
| *Anti-TOX* | PE | REA473 | Miltenyi Biotec | Cat# 130-120-716 |

**Table 2.** Flow cytometry cell dyes.

| Dye Name | Source | Identifier |
|---|---|---|
| *CFSE* | Tonbo | Cat# 13-0850 |
| *Ghost Dye Red 780* | Tonbo | Cat# 13-0865 |
